# Effects of scent enrichment on behavioural and physiological indicators of stress in zoo primates

**DOI:** 10.1101/2020.08.21.260679

**Authors:** Stefano Vaglio, Stefano S. K. Kaburu, Richard Pearce, Luke Bryant, Ailie McAuley, Alexandria Lott, Demi J. Sheppard, Sarah Smith, Beth Tompkins, Emily Elwell, Sara Fontani, Christopher Young, Giovanna Marliani, Pier Attilio Accorsi

## Abstract

Captive breeding is vital for primate conservation, with modern zoos serving a crucial role in breeding populations of threatened species and educating the general public. However, captive populations can experience welfare issues that may also undermine their reproductive success. In order to enhance the well-being of endangered zoo primates, we conducted a study to assess the effects of a new scent enrichment programme on captive red-ruffed lemurs (*Varecia rubra*), black howler monkeys (*Alouatta caraya*), siamangs (*Symphalangus syndactylus*), Lar gibbons (*Hylobates lar*) and orangutans (*Pongo pygmaeus pygmaeus*). We combined behavioural observations and faecal endocrinology analyses to evaluate the effects of a series of essential oils (benzoin, lavender, lemongrass) on five captive troops (N = 19) housed at Dudley Zoo & Castle and Twycross Zoo (UK). We recorded observations of natural species-specific and abnormal stress-related behaviours for 480 hr using instantaneous scan sampling. We collected 189 faecal samples and measured the faecal cortisol concentrations using radioimmunoassay. We found a significant effect of the scent enrichment on behaviours, with red-ruffed lemurs and black howler monkeys reducing their social interactions, as well as red-ruffed lemurs and Lar gibbons decreasing their stress-related behaviours, after they were exposed to the series of essential oils. We also found that red-ruffed lemurs displayed a significant increase in faecal glucocorticoids following the exposure to essential oils. Our contradictory findings suggest that the effects of this series of essential oils may change depending on the species-specific social lives and olfactory repertoires of primates. In conclusion, we cannot recommend using these essential oils widely with zoo primates without additional evaluation.

## Introduction

Almost half of the total primate species recognized today worldwide are classified as endangered or critically endangered in the wild, primarily due to human activities (Estrada et al., 2017). Therefore, raising global scientific and public awareness of the plight of the world’s primates is now vital (Estrada et al., 2017). Zoos may play a major role (Mellor et al., 2015) as zoo animal populations are usually managed to educate the public regarding wildlife and their habitats, and to preserve endangered species through captive breeding and reintroduction programmes (Schulte□Hostedde & Mastromonaco, 2015). However, captive populations, potentially serving as buffers against extinction, experience problems that impair them from being viable for reintroduction into the wild. More specifically, zoo animal populations face reproductive challenges which have so far inhibited them from serving as viable ‘reserve populations’ (Meier, 2016). Additionally, managing zoo populations is challenging because of the mismatch between natural and captive environments and the knock-on effects this has on the repertoire of behaviours exhibited (Carroll et al., 2014). Primates have evolved distinct behavioural patterns, and difficulty in engaging in these behaviours can cause frustration or boredom, which, in turn, can lead to stress and development of abnormal behaviours (Hosey, 2005) that may undermine their individual welfare and ultimately their breeding success.

To maintain captive healthy populations modern zoos take part in conservation breeding programmes. As reproductive success is linked to how closely captive environmental conditions mirror those that primates would be experiencing in the wild (Meier, 2016), zoos also use environmental enrichments to manage captive populations. Environmental enrichments and conservation breeding programmes are directly related, as enrichment is a dynamic, iterative process that changes an animal’s environment, increasing its behavioural choices and prompting a wider range of natural and species-specific behaviours and abilities (Ben-Ari, 2001). Furthermore, enrichment can contribute to promoting resiliency to stress, which helps animals recovering from adverse stimuli (Quirke & O’Riordan, 2011), improving both the exhibit from the visitor perspective and the reproductive performance of the hosted animals (Carlstead & Shepherdson, 1994). Enrichment can also foster the essential skills that animals need for their survival if reintroduced into their habitat (Danial Rioldi, 2013).

Scent-based enrichments can be effective at increasing active behaviours in zoo animals and improve their welfare (Fay & Miller, 2015; Quirke & O’Riordan, 2011; Samuelson et al., 2017). However, this is not always the case and some authors reported findings that are less clear or indicate that scent enrichment has little effect (Wells et al., 2007; Myles & Montrose, 2015). The delivery mechanism of the scent and the type of scents used are crucial for the implementation of novel olfactory enrichment programmes (Baker et al., 2018). The majority of studies have used spices or essential oils rather than focusing on natural or biological scents, but this may not necessarily be appropriate for all species (Wells et al., 2007). The main goal of olfactory enrichment is to improve welfare of animals in captive environments, but there is also the possibility that the use of scents can have additional positive impacts. For example, scents may elicit both behavioural and physiological responses and therefore the use of olfactory enrichment can be potentially used to promote beneficial impacts on reproductive success (Rafacz & Santymire, 2014).

Primates are traditionally considered “microsmatic” (*i*.*e*., with a reduced olfactory sense; Negus, 1958) and, as many uses of enrichment are *ad hoc* and unrecorded, only a small proportion of formal studies on olfactory enrichment has been undertaken on primate species (Clark & King, 2008). However, various lines of evidence suggest that chemical communication may be important in primates (Setchell et al., 2010). In particular, it has become increasingly clear that the sense of smell plays a crucial role in primate socio-sexual communication, with semiochemicals (*i*.*e*., behaviour and physiology-modifying chemicals; Norland & Lewis, 1976) being important for kin recognition, mate choice and the regulation of socio-sexual behaviours (Vaglio et al. 2016). However, little is known about the overall effects of olfactory enrichment on primate species.

The overarching aim of our work is to design and test a new scent enrichment programme to enhance the well-being of critically endangered zoo primates. In this context, we carried out a preliminary study which aimed to assess the effects of a series of essential oils (namely benzoin, lavender and lemongrass) on behavioural and physiological indicators of stress in five captive primate species: red-ruffed lemurs (*Varecia rubra*), black howler monkeys (*Alouatta caraya*), siamangs (*Symphalangus syndactylus*), Lar gibbons (*Hylobates lar*), and orangutans (*Pongo pygmaeus pygmaeus*). As the majority of studies of scent enrichment on zoo primates focus on essential oils, spices or herbs (Wells et al., 2007), we chose three essential oils due to their ecological relevance to non-human primates (benzoin; *e*.*g*., Horvath et al., 2007), effectiveness in domestic animals and humans (lavender; reviewed in Wells, 2009), and efficacy in sheltered cats and dogs as well as in zoo-housed exotic animals (lemongrass; *e*.*g*., Wells, 2004; Ellis & Wells, 2010; Holland, 2018). The primate species investigated in this study are currently classified as critically endangered (red-ruffed lemurs, orangutans), endangered (Lar gibbons, siamangs) or threatened (black howler monkeys) largely due to the deforestation, logging and hunting activities that threaten the habitat and survival of these species across their ranges (IUCN, 2020). Therefore, designing and implementing strategies that improve welfare and breeding success of these species in captivity is particularly crucial.

In this study, we predicted that the scent enrichment would reduce the stress levels of zoo primates, which would be reflected in significant changes in behavioural (*i*.*e*., increase of the frequency of social behaviours, and decrease of the frequency of stress-related behaviours) and physiological (*i*.*e*., decrease of faecal glucocorticoid concentrations, or FGCs) indicators of well-being when comparing before (*i*.*e*., baseline period) and after (*i*.*e*., post enrichment period) the scent enrichment programme. Particularly, this should occur in relatively ‘macrosmatic’ primates (*i*.*e*., primate species with greater levels of olfactory function; Smith & Bhatnagar, 2004) such as lemurs.

## Material and methods

### Study subjects and housing

We studied five captive troops of red-ruffed lemurs, black howler monkeys, siamangs, Lar gibbons, and orangutans housed at Dudley Zoo & Castle (red-ruffed lemurs, Lar gibbons, orangutans) and Twycross Zoo (black howler monkeys, siamangs) in the UK. The troop of red-ruffed lemurs (N = 3) consisted of two related adult males (brothers; both aged 15 years at the beginning of the study period) and one unrelated adult female (aged 14 years). The troop of black howler monkeys (N = 5) was a family group and consisted of one adult female (aged 13 years) and her offspring – one juvenile female (aged 5 years) and three juvenile males (aged 4, 4 and 3 years). The troop of siamangs (N = 3) was a family group and consisted of one adult male (aged 14 years), one adult female (aged 14 years) and their son – one juvenile male (aged 5 years and 6 months). The troop of Lar gibbons (N = 5) was a family group and consisted of one adult male (aged 16 years), one adult female (aged 17 years) and their offspring – one juvenile female (aged 6 years), two young males (aged 2 years and 6 months). The troop of orangutans (N = 3) was a family group and consisted of one adult male (aged 21 years), one adult female (aged 24 years) and their daughter – one juvenile female (aged 5 years).

We carried out behavioural observations and faecal sampling from July to September in 2016, 2017, 2018 and 2019 **(Table 1)**. All troops lived in indoor enclosures (heated to 28 °C) with access to outdoor enclosures (*i*.*e*., “visitor walkthrough” enclosure in the case of red-ruffed lemurs).

**Table 1.** Study protocol. All experimental protocols included a 2-day baseline, 6-day scent enrichment, and 2-day post enrichment.

| Species | Site | Period of time |
| --- | --- | --- |
| Red-ruffed lemurs | Dudley Zoo | July – September 2019 |
| Black howler monkeys | Twycross Zoo | July – September 2016<br>July – September 2019 |
| Siamangs | Twycross Zoo | July – September 2016 |
| Lar gibbons | Dudley Zoo | July – September 2018 |
| Orangutans | Dudley Zoo | July – September 2016<br>July – September 2018<br>July – September 2019 |

### Study protocol

We divided the overall study period into three periods: baseline, scent enrichment, post enrichment. We collected behavioural data and faecal samples for two to six days per study period (10 days in total), two days per week over a 3-month period (one-week baseline; three-week scent enrichment, *i*.*e*. benzoin, lavender, lemongrass; one-week post enrichment), for each species **(Table 1)** in order to use a combination of both behavioural (*e*.*g*., naturalistic species-specific behaviours, stereotypic behaviours) and physiological (*e*.*g*., corticosteroid levels) methods to assess the effects of scent enrichment. *See sections Behavioural Data Collection and Hormone Sampling and Measurements for details*.

#### Scent enrichment

We cut white cotton sheets into 75 cm long and 5 cm wide strips, which were soaked with 20 drops naissance 100% pure essential oil diluted with 12 ml of cold boiled water. We prepared the scent cotton strips during the early morning of each sampling day over the scent enrichment period. We positioned these strips around both indoor and outdoor enclosures; focusing on the outdoor enclosure, we tied them approximately 1 m from the ground around the climbing frames as these were the most used areas of the enclosures. We utilized one essential oil (benzoin, lavender, lemongrass – respectively) per week during the scent enrichment period of the study.

#### Behavioural data collection

We collected behavioural data by instantaneous scan sampling (Altmann, 1974) of some behaviours **(Table 2)**, as a comparable straightforward assessment of major behavioural states which may indicate the expression of significant stress-related (*i*.*e*., self-scratching, pacing) and non-stress-related (*i*.*e*., resting, sleeping, grooming, playing) behaviours, with behaviours recorded at 5-minute intervals over the duration of six hours from 9am to 3pm, ten days over a 3-month period. We recorded a total of 480 hr of observations over the study period, with 50 scan samples each sampling day on each group.

**Table 2.** Ethogram.

| Behaviour | Description |
| --- | --- |
| **Resting** | Lying or sitting while awake, with eyes open and arms down by side of body. |
| **Sleeping** | Lying on back, front or side, eyes closed and whole body is relaxed |
| **Grooming** | Using fingers or mouth to pick through the coat, removing any foreign bodies from a conspecific |
| **Playing** | Animal is engaging in activities such as chasing others, leaping around the enclosure etc. in a playful context |
| **Self-scratching** | An animal rubs their own body at a fast pace |
| **Pacing** | Animal walks back and forth in a distinct, unchanging pattern within the enclosure. |

#### Hormone sampling and measurements

We collected faecal samples every morning before behavioural observations, whenever defecation was observed and the identity of the animal was known. In total, we collected 189 samples (red-ruffed lemurs = 25; black howler monkeys = 56; Lar gibbons = 53; siamangs = 16; orangutans = 39). The samples were stored in a freezer at 20°C right after collection. At the end of the study period, the collated samples were fully prepared by adding biological hazard labels onto each pot before being delivered using dry ice to the Department of Veterinary Medical Sciences and Animal Production Science of Bologna University for radioimmunoassay (RIA).

Cortisol concentrations were determined by RIA. All concentrations were expressed in pg/mg of faecal matter. The extraction methodology followed the methods of Fontani et al. (2014). In brief, five millilitres of a methanol:water (4:1 v/v) solution were added to 60 mg (wet weight) of faeces in capped-glass tube vials. The vials were then vortexed for 30 min using a multitube pulsing vortexer. After centrifugation at 1,500 g for 15 min, 5 ml ethyl ether (BDH Italia, MI, Italy) and 0.2 ml NaHCO3 (5%; Sigma Chemical Co., St. Louis, MO) were added to 1 ml of supernatant. This preparation was vortexed for 1 min and centrifuged for 5 min at 1,500 g. The ether portion was aspirated with a pipette, and evaporated under an airstream suction hood at 37°C. The dry residue was redissolved into 0.5 ml of 0.05 M phosphate-buffered saline (PBS; pH 7.5).

Cortisol was assayed in the faecal samples according to the method of Tamanini et al. (1983). The parameters of the analyses were as follows: sensitivity 3.10 pg/100 l; intra-assay variability 6.8%; interassay variability 9.3%; specificity (%), cortisol 100, corticosterone 9.5, 11,-hydroxyprogesterone 8.3, cortisone 5.3, 11,-desoxycortisol 5.0, progesterone 0.6, desoxycorticosterone 0.5, 20,-dihydrocortisone 0.4, testosterone 0.3, aldosterone 0.1, dehydroepiandrosterone < 0.0001, 5,-pregnenolone <0.0001, 17-estradiol < 0.0001, and cholesterol < 0.0001.

### Statistical analyses

In order to assess the effect of scent enrichment on primate behaviour and FGCs, we first generated three behavioural categories from the individual behavioural measures that we collected. More specifically, we generated: 1) a resting category by adding up our data on resting and sleeping behaviours; 2) a social category by combining our data on grooming and play; and 3) a stress category by combining our data on pacing and self-scratching behaviours(we included scratching in this category since this is commonly considered an indicator of anxiety; Maestripieri et al., 1992). For each behavioural category we ran two sets of analyses: for those species for which we collected data at individual-level (*i*.*e*., black howler monkey, orangutan and siamang) we ran linear mixed model (LMM) analysis, while for those species for which we collected data at group-level (*i*.*e*., red ruffed lemur and Lar gibbon) we used linear regression. For both types of analyses, we included species and enrichment condition (before *versus* after exposure to the scent enrichment) as predictors, while the rates of resting, social and stress-related behaviours were set as dependent variables in separate models. Finally, for the LMM analysis, we set individuals’ ID as random factor. A similar approach was run to test the effect of enrichment condition on FGCs, with the difference that the individual-level LMM analysis included data collected on black howler monkeys, orangutans, siamang *and* red-ruffed lemurs while the regression model was run only on Lar gibbons. For all the analyses, we ran each model twice: one with predictors entered as main effects, and one with predictors entered as interaction. Then, for each analysis, we compared Akaike’s information criterion (AIC) values between the two models in order to find the model with the best fit (*i*.*e*., with the lowest AIC value). Finally, in order to estimate effect size for the LMM models, we use the “r2” function implemented in the “performance” package in R (Nakagawa & Schielzeth, 2013). All models met the assumptions of homogeneity of variance and normality of residuals.

### Ethics Statement

This study followed the guidelines for the care and use of captive animals in the UK, involving non-invasive methods for obtaining both behavioral data and faecal samples from the study subjects. In addition, the study was conducted in compliance with the Convention on International Trade in Endangered Species of Wild Fauna and Flora and approved by the Life Sciences Ethics committee at the University of Wolverhampton (UK) and the Ethics committees at Dudley Zoo & Castle (UK) and Twycross Zoo (UK). We also confirm that our research work was consistent with the American Society of Primatologists’ Principles for Ethical Treatment of Non-Human Primates.

## Results

Our analyses showed that enrichment condition did not have a significant effect on resting rates for any of the species examined **(Table 3)**. Conversely, the LMM analysis examining the effect of scent enrichment on social behaviour among howler monkeys, orangutans and siamangs revealed a significant effect of enrichment condition on rates of social behaviour among these species **(Table 4)**, with eight out of the eleven subjects studied exhibiting a decrease in social behaviour after the introduction of scent enrichment **(Figure 1)**. Only three individuals (two Siamangs and one orangutan) showed an increase in social behaviour after exposure of scent enrichment **(Figure 1)**. Similarly, the regression analysis conducted on Lar gibbons and red-ruffed lemurs showed that the interaction between species and enrichment condition had a significant impact on rates of social behaviour **(Table 4)**. This analysis revealed that while rates of social interactions among Lar gibbons were comparable between before and after exposure to the scent enrichment, among red ruffed lemurs the introduction of scent enrichment significantly reduced rates of social behaviour **(Figure 2)**.

**Table 3.**
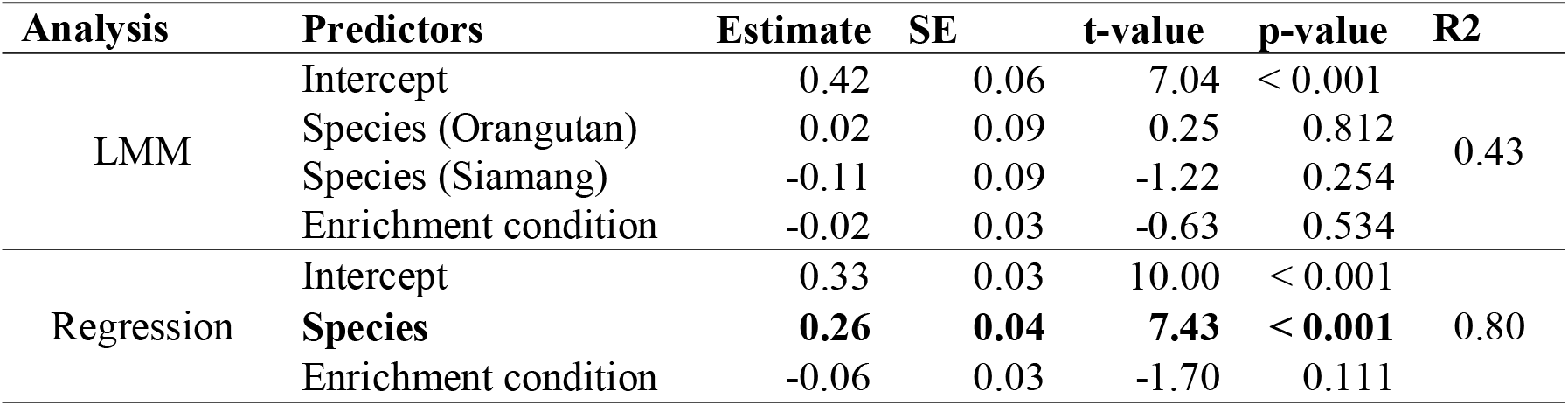
Results of LMM and regression analyses testing the effect of Enrichment condition, and Species on resting rates. Significant result is shown in bold.

**Table 4.** Results of LMM and regression analyses testing the effect of Enrichment condition, and Species on social rates. Significant results are shown in bold.

| Analysis | Predictors | Estimate | SE | t-value | p-value | R2 |
| --- | --- | --- | --- | --- | --- | --- |
| LMM | Intercept | 0.08 | 0.03 | 2.65 | 0.028 | 0.18 |
|  | Species (Orangutan) | 0.02 | 0.04 | 0.43 | 0.679 |  |
|  | Species (Siamang) | -0.02 | 0.04 | -0.42 | 0.683 |  |
|  | **Enrichment condition** | **0.04** | **0.02** | **2.05** | **0.043** |  |
| Regression | Intercept | 0.14 | 0.02 | 9.38 | < 0.001 | 0.46 |
|  | **Species** | **-0.06** | **0.02** | **-2.75** | **0.017** |  |
|  | Enrichment condition | -0.02 | 0.02 | -0.86 | 0.405 |  |
|  | **Enrichment condition \* Species** | **0.07** | **0.03** | **2.54** | **0.025** |  |

**Figure 1.**
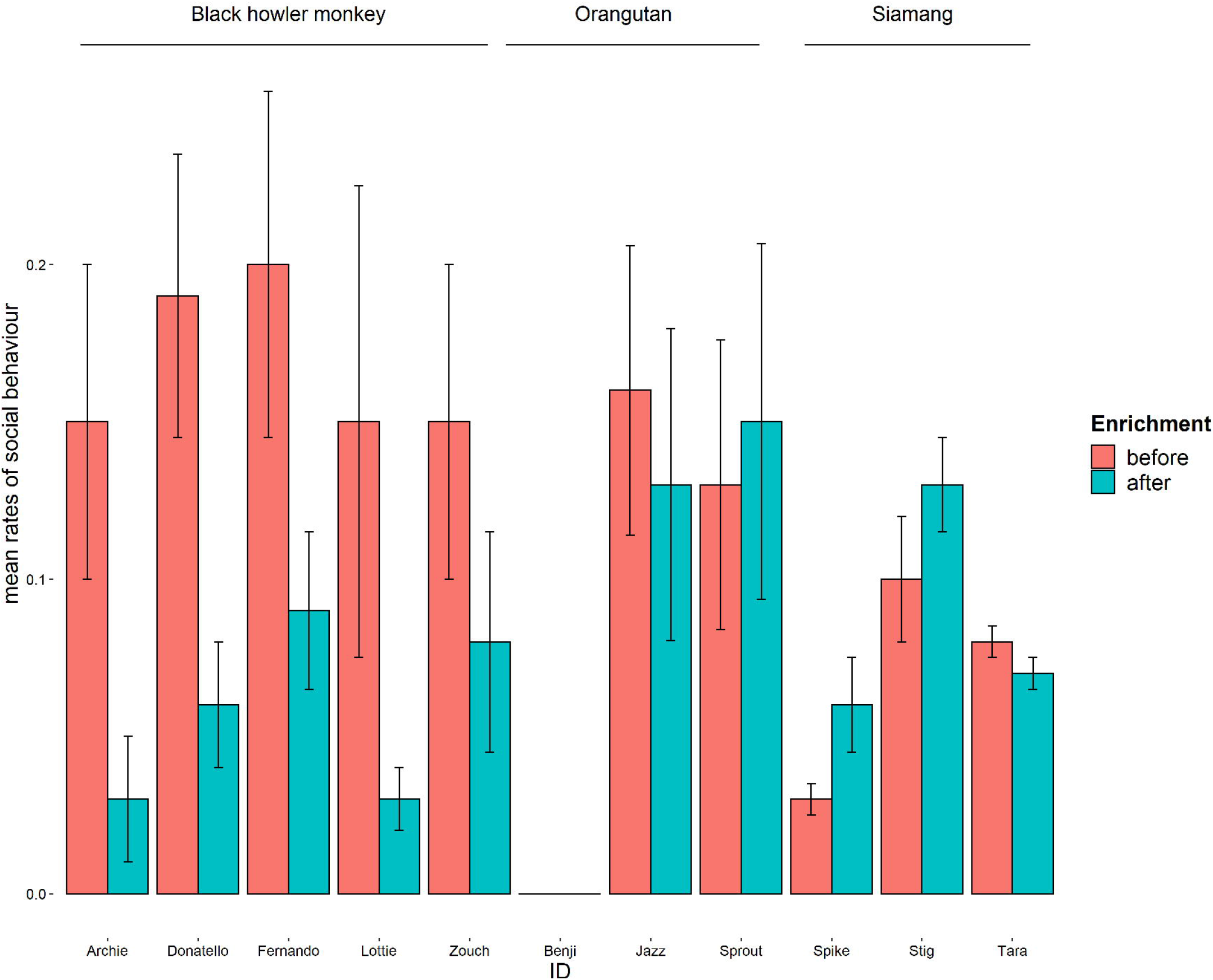
Mean rates (± SEM) of social behaviour per individual among black howler monkey, orangutan and siamang. The LMM analysis revealed a significant effect of the scent enrichment on social behaviours, with eight out of the eleven subjects studied exhibiting a decrease in social interactions after the introduction of the essential oils.

**Figure 2.**
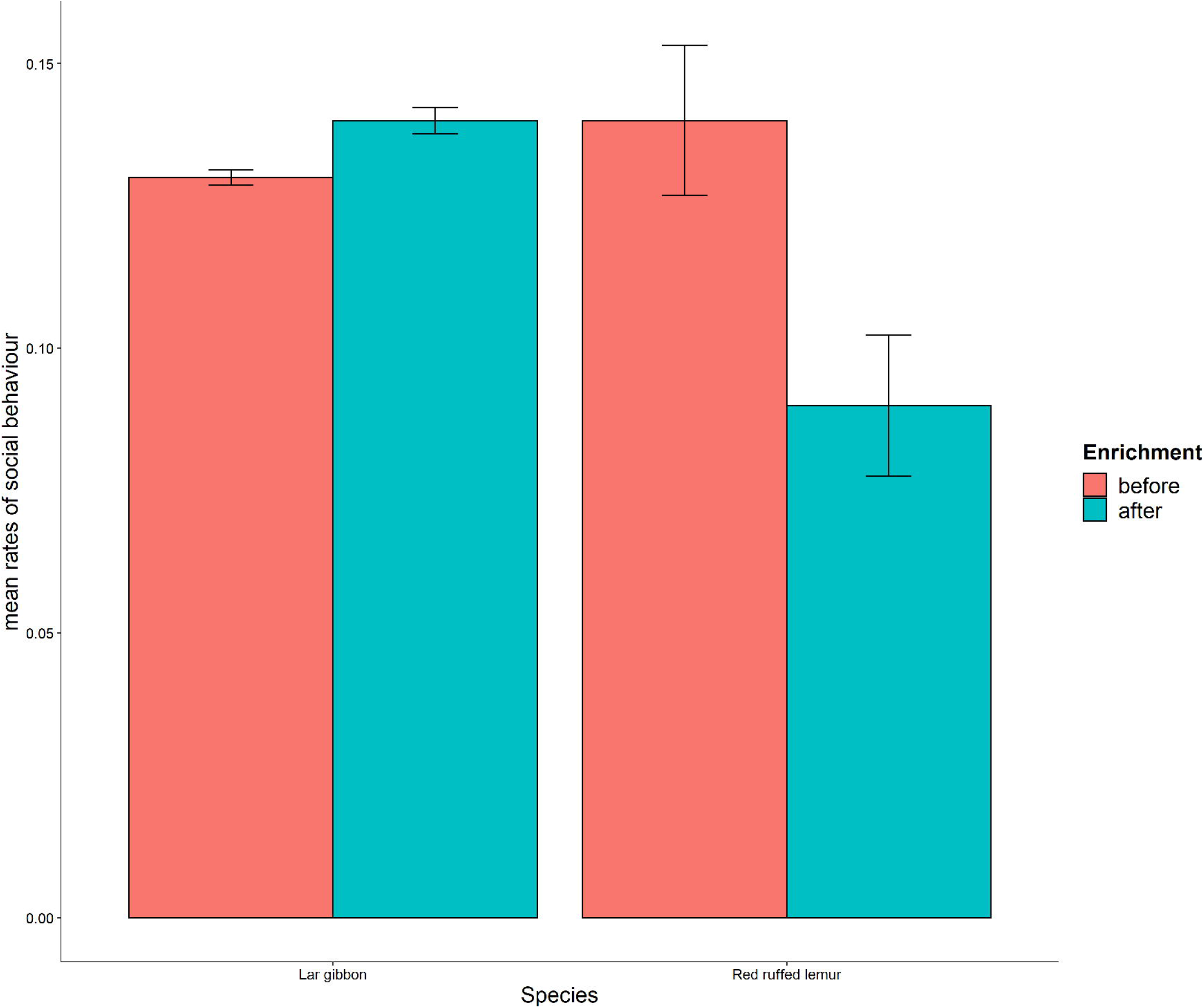
Mean rates (± SEM) of social behaviour among Lar gibbon and red-ruffed lemur. The regression analysis showed that the introduction of the scent enrichment induced a significant reduction in social behaviours among red-ruffed lemurs.

We did not find any significant effect of enrichment condition on rates of stress-related behaviour on howler monkeys, orangutans and siamangs via the LMM analysis **(Table 5)**. By contrast, we found a significant effect of the enrichment condition on rates of stress-related behaviour among Lar gibbons and red-ruffed lemurs in the regression model. This analysis showed that both species exhibited a significant reduction in rates of stress-related behaviour following the exposure to scent enrichment **(Figure 3)**.

**Table 5.** Results of LMM and regression analyses testing the effect of Enrichment condition, and Species on rates of stress-related behaviour. Significant results are shown in bold.

| Analysis | Predictors | Estimate | SE | t-value | p-value | R2 |
| --- | --- | --- | --- | --- | --- | --- |
| LMM | Intercept | -0.002 | 0.03 | -0.06 | 0.96 | 0.78 |
|  | Species (Orangutan) | 0.003 | 0.04 | 0.09 | 0.93 |  |
|  | Species (Siamang) | 0.104 | 0.04 | 2.49 | 0.04 |  |
|  | Enrichment condition | 0.007 | 0.01 | 1.00 | 0.32 |  |
| Regression | Intercept | 0.01 | 0.01 | 1.37 | 0.192 | 0.30 |
|  | Species | 0.00 | 0.01 | -0.49 | 0.633 |  |
|  | **Enrichment condition** | **0.02** | **0.01** | **2.42** | **0.030** |  |

**Figure 3.**
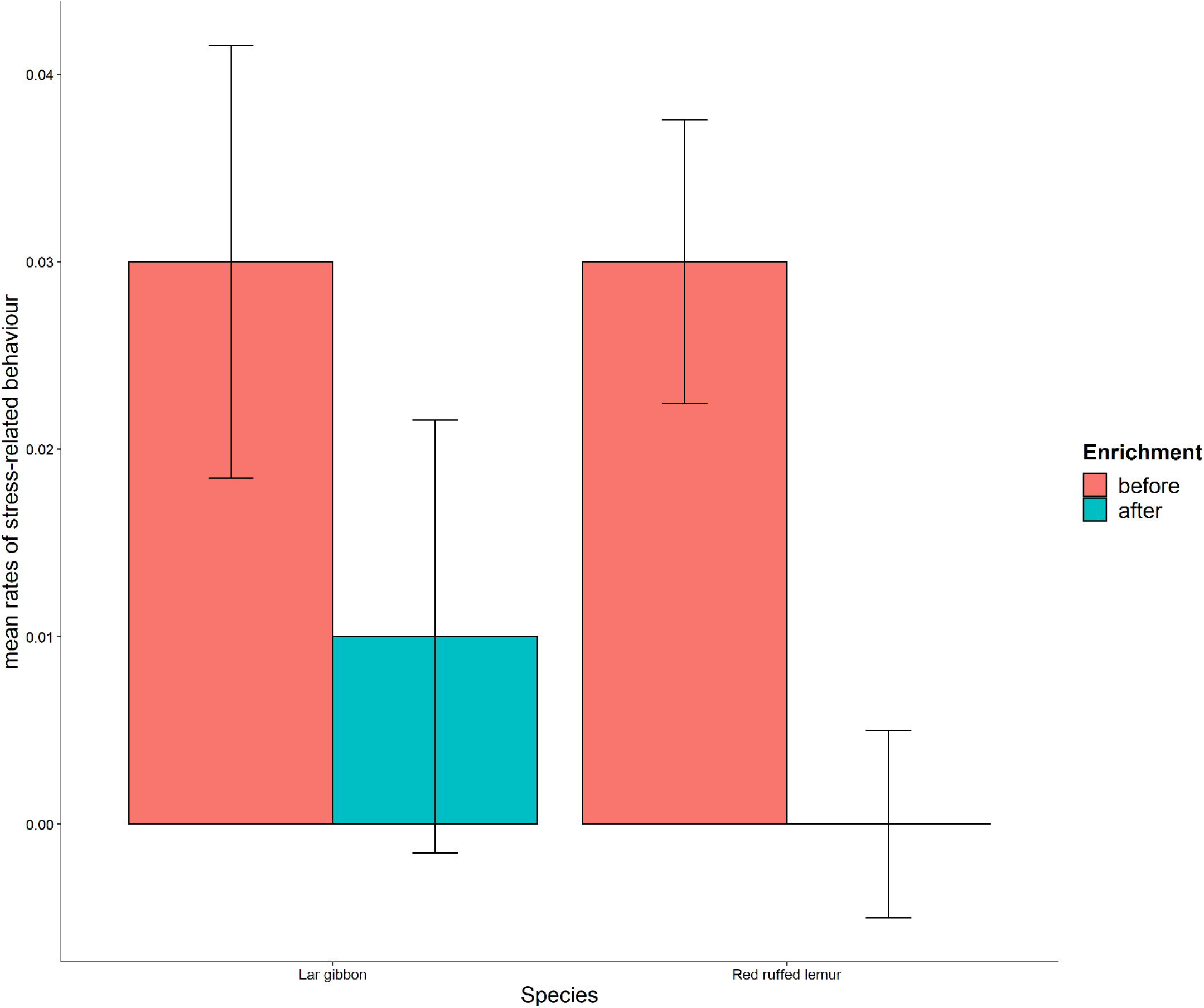
Mean rates (± SEM) of stress-related behaviours among Lar gibbon and red-ruffed lemur. The regression model showed that both Lar gibbon and red-ruffed lemur exhibited a significant reduction in stress-related behaviours following the exposure to the scent enrichment.

The LMM model that investigated the effect of scent enrichment on FGCs among howler monkeys, orangutans, siamangs and red-ruffed lemurs revealed a significant interaction between species and enrichment condition **(Table 6)**. More specifically, the analysis showed that enrichment condition affected FGCs in red-ruffed lemurs but not in other study species. Contrary to our expectations, however, we found that FGCs increased after exposure to scent enrichment, compared to before the introduction of the scent **(Figure 4)**. Interestingly, **Figure 4** shows that orangutans seemed to decrease their FGC levels following exposure to scent enrichment, although the effect failed to reach statistical significance. Finally, among Lar gibbons, although mean FGC concentrations increased after the introduction of scent enrichment **(Figure 4)**, the regression analysis did not reveal any significant effect of enrichment condition on FGCs (Estimate = −1.71, SE = 1.82, t-value = −0.94, p = 0.35, R2 = 0.04).

**Table 6.** Results of LMM analysis testing the effect of Enrichment condition, and Species on FGCs.

| Analysis | Predictors | Estimate | SE | t-value | p-value | R2 |
| --- | --- | --- | --- | --- | --- | --- |
| LMM | Intercept | 0.91 | 0.40 | 2.28 | 0.028 | 0.28 |
|  | Species (Orangutan) | -0.46 | 0.65 | -0.70 | 0.491 |  |
|  | Species (Red ruffed lemur) | 1.30 | 0.84 | 1.55 | 0.130 |  |
|  | Species (Siamang) | -0.87 | 0.76 | -1.14 | 0.263 |  |
|  | Enrichment condition | 0.27 | 0.59 | 0.46 | 0.648 |  |
|  | Enrichment condition * Species (Orangutan) | 1.25 | 0.83 | 1.49 | 0.141 |  |
|  | **Enrichment condition \* Species (Red ruffed lemur)** | **-2.04** | **0.95** | **-2.16** | **0.035** |  |
|  | Enrichment condition * Species (Siamang) | -0.26 | 1.27 | -0.21 | 0.838 |  |

**Figure 4.**
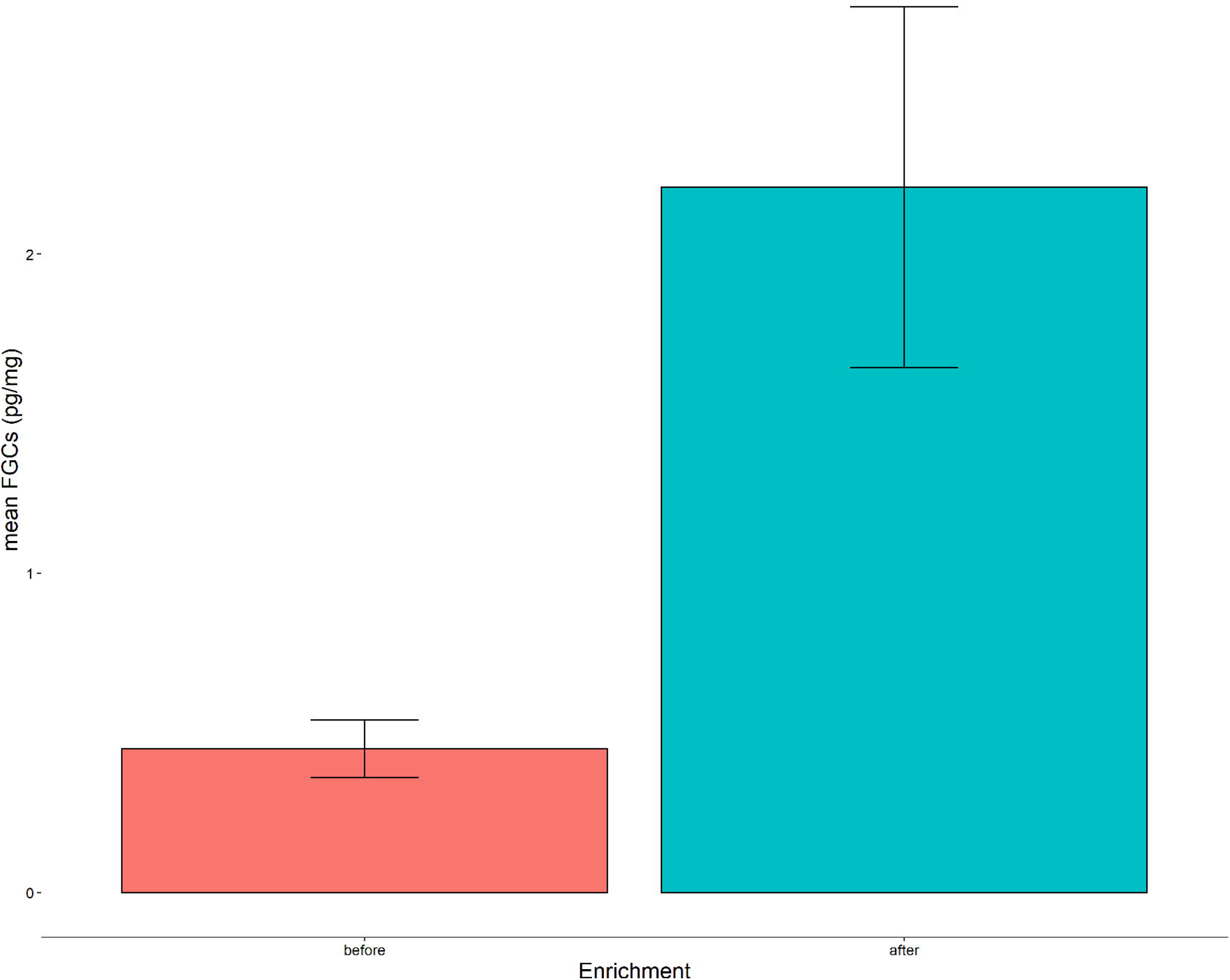
The LMM model showed that the scent enrichment elicited an increase in FGC levels in red-ruffed lemurs, but not in any other study species.

## Discussion

The effects of scent enrichment have previously been tested on several domestic, farm, laboratory and zoo-housed animals (Blackie & de Sousa, 2019; Heitman et al., 2018). However, olfactory stimulation is still one of the least studied forms of enrichment (reviewed in Campbell-Palmer & Rosell, 2011). In addition, there are mixed and conflicting assumptions regarding the benefits of olfactory enrichment on animal welfare, and this is particularly the case of primate species, among which the overall effects of scent enrichment are still unclear and understudied (reviewed in Wells, 2009).

Unexpectedly we found a significant reduction in rates of social interactions after being exposed to the series of essential oils in both red-ruffed lemurs and black howler monkeys. By contrast, several authors have found that scent enrichment may cause increasing active behaviours in zoo-housed non-primate species, such as cheetahs (*Acinonyx jubatus*) (Quirke & O’Riordan, 2011), Californian sea lions (*Zalophus californianus*) (Samuelson et al., 2017) and Rothschild giraffes (*Giraffa camelopardalis rothschildi*) (Fay & Miller, 2015), but not in meerkats (*Suricata suricatta*) (Myles & Montrose, 2015). Regarding primates, Gronqvist et al. (2013) showed that olfactory enrichment significantly increased the frequency of natural species-specific behaviours in captive Javan gibbons (*Hylobates moloch*) although the interest in the new scent decreased rapidly after the first day, whilst no significant effects on individual behaviours were found in ring-tailed lemurs (*Lemur catta*) (Baker et al., 2018) and gorillas (*Gorilla gorilla gorilla*) (Wells et al., 2007). The effect that our scent enrichment exerted on social behaviours, with decreased rates of social interactions in red-ruffed lemurs and black howler monkeys, but no significant effects on siamangs, Lar gibbons and orangutans, might be related to differences in social organisations and structures among these species. Specifically, red-ruffed lemurs and black howler monkeys are social species living in small groups including both adult males and females, while siamangs and Lar gibbons are monogamous and orangutans are solitary. Red-ruffed lemurs and black howler monkeys, thus, display more social affiliative behaviours which have a stress-reducing effects. We, therefore, speculate that red-ruffed lemurs and black howler monkeys could have reduced their rates of social behaviours because our scent enrichment might have decreased the need for reassurance-derived social interactions. However, we recognize that further factors may have induced such differences between the effects of our scent enrichment on individual species; for instance, it is possible that the new unfamiliar scents increased the stress levels in red-ruffed lemurs because they perceived them more intensely than the other study species, or that decreased rates in social behaviours in red-ruffed lemurs and black howler monkeys are due to increased rates in other behaviours (such as inspections, locomotion, etc.) which were not measured during our study.

We also found a significant reduction in rates of stress-related behaviours after red-ruffed lemurs and Lar gibbons were exposed to the series of essential oils, which is the most promising outcome of this preliminary study about the potential positive effect of such essential oils. Similar findings have been reported in non-primate species. For example, Uccheddu et al. (2018) exposed domestic dogs to a variety of essentials oils and found that some scents increased frequencies of behavioural indicators of relaxation while others decreased behavioural indicators of stress, such as pacing and over-grooming. Similarly, a study on cheetahs and Sumatran tigers (*Panthera tigris sumatrae*) found that stereotypic pacing behaviour significantly decreased in the presence of a hay ball with cinnamon (Damasceno et al., 2017). The significant effect from our series of essential oils on stress-related behaviours in red-ruffed lemurs and Lar gibbons is consistent with our prediction that scent enrichment would reduce behavioural indicators of stress; however, we acknowledge the conflicting findings related to the lack of effectiveness of our enrichment shown in siamangs, Lar gibbons and orangutans.

Our finding that red-ruffed lemurs showed a significant increase in FGC levels following the exposure to the series of essential oils may suggest different interpretations. First, this result implies that changes in behavioural indicators of stress, such as pacing and self-scratching, do not necessarily mirror changes in physiological indicators of stress, such as FGCs. This is consistent with the study by Higham et al. (2009) on olive baboons (*Papio anubis*) showing that day-to-day variation in FGC concentrations do not correlate with changes in self-directed behaviours and suggesting that these indicators may signpost two different types of stress, with self-directed behaviours reflecting low-level acute stress or anxiety (Maestripieri et al., 1992) while FGCs reflecting high-level chronic stress (Sapolsky, 1992). Accordingly, self-directed behaviours have been found to increase in anxiety-inducing contexts, such as when animals are given anxiogenic drugs (Schino et al., 1996) or after aggression (Schino, 1998); FGC concentrations have been shown to increase when animals are exposed to high levels of stress, such as in the presence of tourists (Barja et al., 2007) or when exposed to the odour of a predator (Monclús et al., 2006). Additionally, although glucocorticoids are commonly associated with the negative aspects of stress, these steroid hormones play many important roles both in mediating the response to stress and in the circadian rhythm (McEwen, 2019). Thus, another potential explanation for elevated FGCs is increased energy expenditure. For instance, it is possible that increased FGCs in red-ruffed lemurs may be due to enhanced positive arousal related to increased rates in other behaviours (such as investigatory behaviors and locomotion) which we did not measure during our study. Hence, as suggested by other authors (reviewed by Hosey et al., 2013), we emphasize that *both* behavioural *and* physiological indicators should be used to investigate the stress levels of individual animals, while behavioural indicators of anxiety alone should not be interpreted as definite indicators of glucocorticoid production. Interestingly, we found that our scent enrichment exerted both behavioural and physiological effects only on red-ruffed lemurs. Although primates have traditionally been considered to be “microsmatic” with a simultaneous amplified emphasis on vision (Dominy & Lucas, 2001; Fornalé et al., 2012; Gerald, 2003), several studies suggest that chemical communication is important also for primate species (reviewed by Drea, 2020). Particularly, it is established that some species rely heavily on olfaction in addition to vision and auditory senses; for instance, this is the case of several lemurs (Gould & Overdorff, 2002; Janda et al., 2019; Scordato & Drea, 2007) and squirrel monkeys (Laska et al., 2000). This would explain the significant impact of our scent enrichment on red-ruffed lemurs, rather than the other study species for which no such response was observed, as lemurs have retained a greater olfactory complexity than other lineages such as monkeys and apes. However, we recognize that other factors may have contributed to such effects of our scent enrichment on individual species; for instance, it is possible that the effect on red-ruffed lemurs could be related to their different enclosure design (*i*.*e*., “visitor walkthrough” – including a section in which the public could be very close) which ultimately could have led the lemurs being exposed to a different olfactory environment (*i*.*e*., anthropogenic) than the other study species.

Finally, we have to acknowledge some major limitations of this preliminary study. First of all, although our study is ambitious in many respects (*i*.*e*., we worked on several species, over several years, across three conditions and with multiple measures intended to assess welfare), we focused on limited data pools which included relatively small sample size and unit of analysis. Additionally, we did not record behaviors, such as normal locomotion, foraging, inspections and investigatory actions (*e*.*g*., exploring around the scented cloths), but changes in these behaviors could also be very informative.

## Conclusion

This preliminary study provided contradictory findings and suggested that the application of our new scent enrichment programme may affect the stress levels of zoo-housed primates; particularly in the case of primate species where odour plays a crucial role, such as red-ruffed lemurs. Following the exposure to the series of essential oils (benzoin, lavender, lemongrass), both red-ruffed lemurs and Lar gibbons exhibited significantly lower rates of stress-related behaviours, such as pacing and self-scratching. Conversely, red-ruffed lemurs also significantly increased their levels of FGCs, which however might be explained by an increase in positive arousal. However, given that the exposure to the series of essential oils entailed a significant reduction in social behaviours in red-ruffed lemurs and black howler monkeys as well as a significant increase in FGCs in red-ruffed lemurs, we cannot even exclude negative effects by our scent enrichment. Therefore, in conclusion, we cannot recommend using this series of essential oils widely without further evaluation.

Future work would need to expand the investigation of the effect of our scent enrichment on primate welfare by focusing on both a larger sample size and a wider range of species across the major lineages. Also, it would be crucial to test further types of scent enrichment by considering the ecological/biological relevance of the scent enrichment to the study species. Many scents, including essential oils, are chosen based on their effectiveness in humans or domestic animals, but this may not necessarily be appropriate for all animal species (Wells, 2009). In particular, as previous authors have suggested, important factors to consider for the implementation of novel olfactory enrichment programmes are the mechanism of delivery of the scent and the type of scents used (Baker et al., 2018), while the effectiveness of any intervention should be continually monitored to inform best practices.

## Acknowledgements

We are grateful to Dudley Zoo & Castle (especially David Beeston, Chris Leeson, Pat Stevens, and primate keepers) and Twycross Zoo (especially Mat Liptovszky, Manuela Townsend, Freisha Patel, Jessica Rendle, and primate keepers) for their support to the project and assistance with sample collection. We thank Jemma Billingsley, Stephanie Courten and Henry Swain for collecting additional data which provided helpful insight but could not be included in this study. Finally, we thank two anonymous reviewers for their constructive comments and suggestions. This research work was supported by the Faculty of Science and Engineering, University of Wolverhampton (equipment & laboratory consumables), and the Department of Veterinary Medical Sciences, University of Bologna (laboratory analyses). This project also received funding from the Primate Society of Great Britain (Captive Care Grant – round 2018 to S.V.).

## Author contribution

SV conceived and designed the study. SS, BT, AM, AL, AQ, DJS, LB and RP collected and organised the behavioural data and faecal samples. GM and PAA measured the hormone levels. SSKK analysed the data. SV, SSKK, CY, EE and SF equally contributed to writing the paper.

## Data availability statement

The raw data are available on request.

